# Fire traps in the wet subtropics: new perspectives from Hong Kong

**DOI:** 10.1101/2023.11.08.566152

**Authors:** Aland H. Y. Chan, David A. Coomes

## Abstract

1. Fires undermine efforts to restore degraded forests in the wet tropics and subtropics. Grasslands and shrublands established after fires are more fire-susceptible than forests and tend to be set alight more often, creating a positive feedback loop that curbs succession (i.e. create a fire trap). Understanding the factors that underpin the strength of these fire traps could transform restoration programmes by identifying the steps needed to escape them.
2. Fire traps are notoriously challenging to quantify because multiple factors influence fire occurrence and vegetation recovery. Here we used multi-decadal satellite imagery from Landsat to create a 34-year time series of burn areas and vegetation dynamics in wet subtropical Hong Kong. These dynamic maps were then used to characterise (1) the influence of successional stage on fire occurrence, having accounted for topographical and ignition source imbalances using neighbourhood analyses and entropy balancing (EBAL) weights, and (2) recovery time to the next successional stage by survival analysis with EBAL weights.
3. Our analyses revealed that fire regimes in the wet subtropics are defined by strong fire-vegetation feedbacks. Grasslands and shrublands were 20 and 9 times more susceptible to fires than forests in similar topographic positions. Human activities compounded these differences by disproportionally introducing more ignition sources to grasslands (2.3 times) and shrublands (2.0 times) than to forests.
4. Burnt shrublands recovered to forests faster (19 years) than grasslands (40 years). Proximity to forest patches had strong positive effects on recovery rates, highlighting the importance of seed sources. Post-fire recovery was faster on wetter northwest-facing sites and valleys. Overall, topography strongly influenced recovery processes but hardly affected fires occurrence.
5. *Synthesis and applications*. Our study provides the first quantification of fire-trap processes in the wet subtropics, which provides new opportunities for evidence-based fire suppression and post-fire restoration. Our results suggest that (1) fire traps could be mitigated by fire-suppression programmes as the traps are currently exacerbated by ignition source imbalance; (2) establishing green fire breaks represent an effective fire suppression measure in the wet subtropics; and (3) active restoration could target areas where models predict sluggish post-fire natural regeneration.

## 1. Introduction

Humans have fundamentally changed fire regimes in the wet tropics and subtropics. Naturally, pristine rainforests retain moisture well. Barring extreme droughts, these forests are naturally fire-resistant with fire return intervals of 100-1000 years (Goldammer and Seibert, 1989; Goldammer, 1990; Cochrane, 2003). However, up to 30-40% of all tropical forests are now degraded by logging and agricultural activities (Budiharta *et al*., 2014). Dominated by C4 grasses, *Dicranopteris* fern mats, short bamboos, and shrublands, degraded wet tropical landscapes retains moisture poorly and are much more likely to burn during dry spells (Matos, Santos and Chevalier, 2002; Haberle *et al*., 2010; Hoffmann, Geiger, *et al*., 2012; Hoffmann, Jaconis, *et al*., 2012). Fires, in turn, disproportionately kill saplings of fire-sensitive late-successional tree species. Meanwhile, grasses, forbs and shrubs tend to have basal meristems, lignotubers, or other forms of underground energy stores (De Moraes *et al*., 2016; Paula *et al*., 2016; Simpson *et al*., 2016). These features help plants survive by exploiting the steep temperature gradients created by soil insulation (Beadle, 1940). By resprouting and dispersing into burnt patches, grasses and shrubs reinforce its dominance in fire-disturbed habitats (Paula *et al*., 2016; Simpson *et al*., 2016). In many degraded landscapes, this leads to positive fire-vegetation feedbacks, which may create "fire traps" that perpetuate early-successional vegetation in areas where the climate supports closed-canopy forests (Bell, 1984; Hoffmann, Geiger, *et al*., 2012; Flores *et al*., 2016; Staal *et al*., 2018; Van Nes *et al*., 2018; Mata *et al*., 2022). These effects are further compounded by the abundance of anthropogenic ignition sources (Cochrane, 2003; Tien Bui *et al*., 2016) and stronger droughts under climate change (Hoffmann, Schroeder and Jackson, 2003; Seidl *et al*., 2017; Lizundia-Loiola, Pettinari and Chuvieco, 2020; Clarke *et al*., 2022).

Properly describing and quantifying fire-trap dynamics in the wet tropics is crucial for the management and restoration of degraded wet tropical landscapes. International initiatives, such as the Bonn Challenge, UN Decade of Ecosystem Restoration, and the One Trillion Tree initiative, have repeatedly called for large-scale restoration in the wet tropics (Lamb, Erskine and Parrotta, 2005; Secretariat, 2010; Verdone and Seidl, 2017). Understanding fire-trap dynamics in degraded wet tropical and subtropical sites is critical for developing viable, cost-effective, and climate-resilient local restoration strategies (Scheper, Verweij and van Kuijk, 2021). Assessing the relative importance of vegetation structure, anthropogenic ignition source distribution, and local topography on fire occurrence is the first step towards targeted fire suppression (Carmo *et al*., 2011). Similarly, evaluating how post-fire recovery rate is affected by pre-fire vegetation structure, burn severity, and topographical factors helps land managers decide where and when intervention is needed (Souza-Alonso *et al*., 2022).

Fire susceptibility and post-fire recovery in the wet tropics and subtropics are currently understudied. Despite evidence showing that degraded sites in the wet tropics are reasonably fire-prone (Uhl, Kauffman and Cummings, 1988; Flores *et al*., 2016; Mata *et al*., 2022), there is still a prevailing perception of these bioregions being "too wet to burn". Existing studies on fire dynamics continue to focus on naturally fire-adapted Mediterranean, boreal, and savanna ecosystems (Mallek *et al*., 2013; van Butsic, Kelly and Moritz, 2015; Kibler *et al*., 2019; Qiu *et al*., 2021; Kurbanov *et al*., 2022). Since wet tropical landscapes are rarely fuel-limited and have distinctive fire-vegetation dynamics (Tepley *et al*., 2018), it is questionable whether patterns observed in other ecosystems hold true in wet tropical and subtropical biomes. Existing studies also does not fully address the issue of covariate imbalance when evaluating fire susceptibility amongst different vegetation types. For instance, it is reasonable to expect forests to disproportionally occupy wetter valleys, while grasslands dominate the drier ridgetops. This is further complicated by the non-random distribution of ignition sources (Oliveira *et al*., 2012; Tien Bui *et al*., 2016). Grasslands and shrublands could be more exposed to ignition sources as they are closer to settlements and more accessible to humans. To accurately quantify fire traps, these relationships need to be carefully untangled. Similarly, post-fire recovery trajectories in the wet tropics are worth re-evaluating as they determine whether sites could escape the fire trap. Previous studies have identified fire frequency, distance from forest patches, burn severity, soil type, species composition, and several topographical variables as factors that affect rate of post-fire recovery (Ireland and Petropoulos, 2015; Goosem *et al*., 2016; Araújo *et al*., 2017; Bright *et al*., 2019; Rochimi, Aj and Meador, 2021; Kurbanov *et al*., 2022; Marsh *et al*., 2022), but a systematic evaluation of the importance of these variables in the wet tropics is currently lacking. Crucially, many existing studies quantified post-fire recovery by tracking the rebound of remotely-sensed indices, such as the normalized difference vegetation index (NDVI) or normalized burn ratio (NBR), to pre-disturbance values (Gouveia, DaCamara and Trigo, 2010; Ireland and Petropoulos, 2015; Fernández-García *et al*., 2018; Bright *et al*., 2019; Pérez-Cabello, Montorio and Alves, 2021; Kurbanov *et al*., 2022). In the degraded wet tropics, however, the background landscape is itself on a succession trajectory. Burnt areas rapidly revegetate and return to pre-fire conditions quickly (Idris, Kuraji and Suzuki, 2004; Melchiorre and Boschetti, 2018; Chan *et al*., 2023), but returning the system to the pre-fire degraded condition is often not the goal of land managers. Instead, it is more appropriate to study recovery time to forests after fire, yet none of the existing studies have adopted such an approach to analyse post-fire recovery over large scales (Kurbanov *et al*., 2022).

Methodological advances in remote sensing and biostatistics have made it increasingly manageable to track fire susceptibility and post-fire recovery across large spatiotemporal scales. Accurate burn area mapping in the wet tropics and subtropics is technically challenging (Alencar *et al*., 2022; Chan *et al*., 2023). High cloud cover and rapid burn area revegetation have largely limited full burn area mapping across decadal time scales to satellites with short return times (e.g. MODIS), which led to tradeoffs in accuracy and ground resolution (Humber *et al*., 2019; Szpakowski and Jensen, 2019; Franquesa *et al*., 2022; Chan *et al*., 2023). Advances in imagery processing have, however, lifted many of these restrictions. Long-term Landsat-based burn area products, which have 30 m ground resolutions and are better at detecting small burnt patches than MODIS-based maps, are now available for parts of the wet tropics and subtropics (Alencar *et al*., 2022; Chan *et al*., 2023). Additionally, new statistical tools have also made it easier to handle data extracted from burn area and vegetation time series. There are now standardised workflows to handle covariate imbalance, such as the tendency for forests to be found in wetter valleys, by matching or reweighting (Cannas and Arpino, 2019; Markoulidakis *et al*., 2022). Survival analysis, which is used to model fire-vegetation feedbacks (Reed *et al*., 1998; Tepley *et al*., 2018) and vegetation succession (Longpre and Morris, 2012), has also developed rapidly. The integration with machine learning to create random survival forests now make it possible to easily visualise variable importance and make survival time predictions, especially for large datasets with non-linear or covarying predictors of survival (Ishwaran *et al*., 2008).

In this study, we utilised these remote sensing and statistical approaches to (1) quantify the strength of fire-vegetation feedbacks relative to other predictors of fire occurrence (e.g. topographical position) and (2) describe how different factors affect post-fire recovery rates. We conducted the study in the extensive wet subtropical landscapes of Hong Kong, which was heavily degraded in the past and experiences a large number of anthropogenic fires despite restoration efforts. The objective is to use the results to identify effective approaches for fire suppression and active restoration of burnt areas.

## 2. Methods

### 2.1 Study area

The study was conducted in the wet subtropical countryside of Hong Kong (22° 16’ 8’’ N, 113° 57’ 6’’E) (**Figure 1**). On average, the region receives over 2400 mm of rainfall per year and would have historically been covered by evergreen broadleaved subtropical rainforests (Dudgeon and Corlett, 2004; Abbas, Nichol and Fischer, 2016; Yang *et al*., 2018). However, centuries of human settlement and agricultural activity had decimated >90% of the natural forests, creating a barren landscape of grasslands and short shrublands (Zhuang and Corlett, 1997; Dudgeon and Corlett, 2004). After the second world war, an economic transition led to a sharp fall in the rural population and associated land management practices (Hau *et al*., 2005). Widespread agricultural abandonment, along with the designation of strictly protected Country Parks over 40% of the land area, led to over 70 years of natural succession and assisted regeneration (Abbas, Nichol and Fischer, 2016). The current vegetation of Hong Kong consists of a mosaic of grasslands, shrublands, secondary forests, and plantations (Kwong *et al*., 2022).

**Figure 1:**
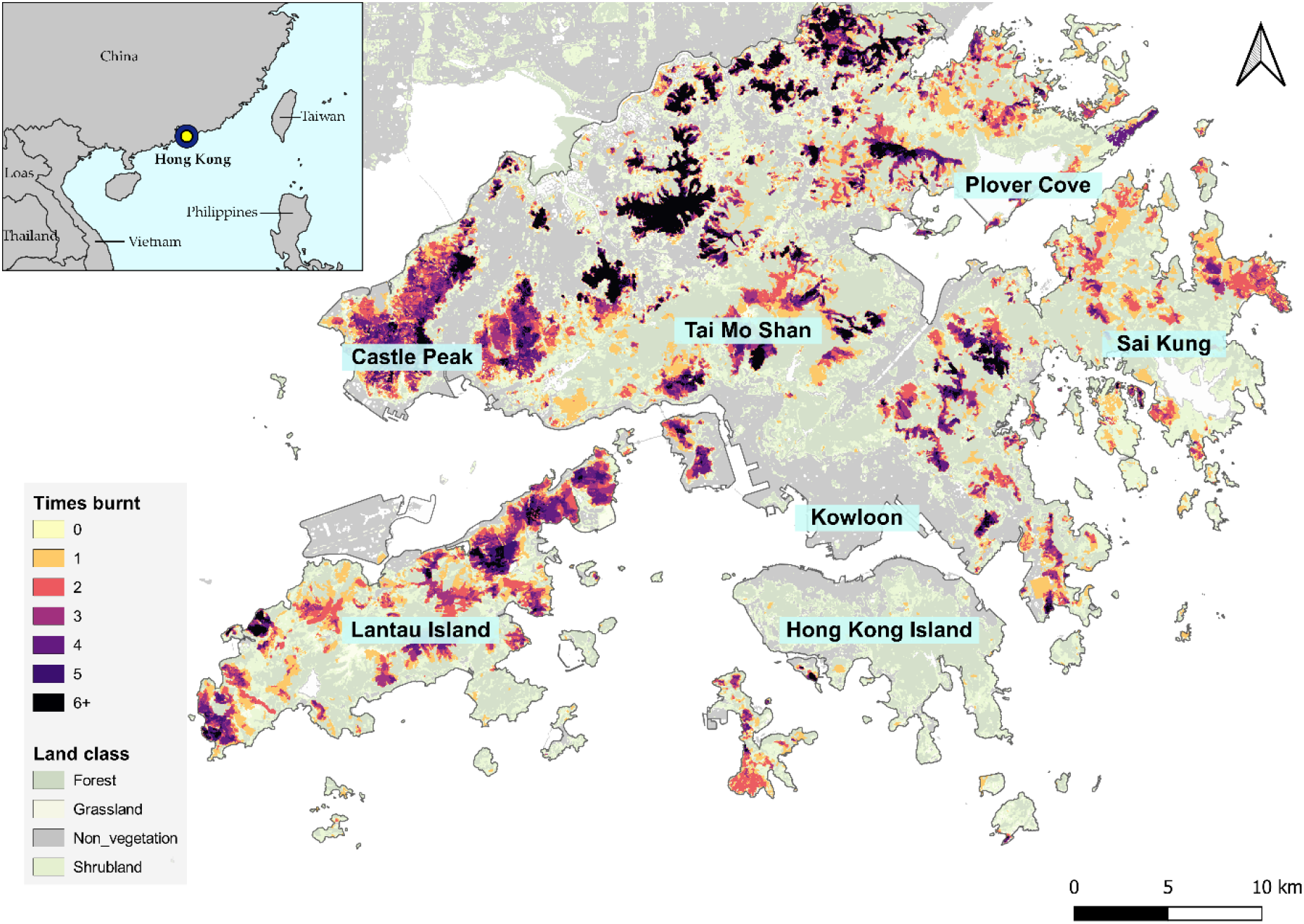
The study area of Hong Kong. Burnt areas detected by the LTSfire pipeline between 1986 and 2020 are overlaid on a Landsat based land cover map of 2013-2014 (Chan et al., 2023).

Natural fires are rare in Hong Kong as the main natural ignition source (lightning) is usually accompanied by torrential rain, but anthropogenic fires are frequent (Fung and Jim, 1993; Dudgeon and Corlett, 2004; Chan *et al*., 2023). Common anthropogenic ignition sources include joss paper burnt around graves in local festivals, cigarette butts, and campfires (Chau, 1994; Chan, 2005). The Fire Services Department received 4561 reports of vegetation fires between 2016 and 2020. Overall, 287 km^2^ (39%) of the 728 km^2^ of vegetated area burnt at least once between 1986 and 2020, with 65% of the affected area burning more than once (Chan *et al*., 2023). The diverse vegetation structure coupled with notable fire occurrence creates a convenient setting to study fire traps.

### 2.2 Overview of methods

Overall, we used (1) Landsat imagery to create burn area and vegetation maps for the study period (1986 – 2020) and (2) LiDAR data to calculate topographical variables. We then used these products to study how different factors affected fire occurrence, fire susceptibility, and post-fire recovery rates. **Figure 2** provides a methodological flow chart with numbers serving as a guide to the relevant sections where the detailed methodologies are described.

**Figure 2:**
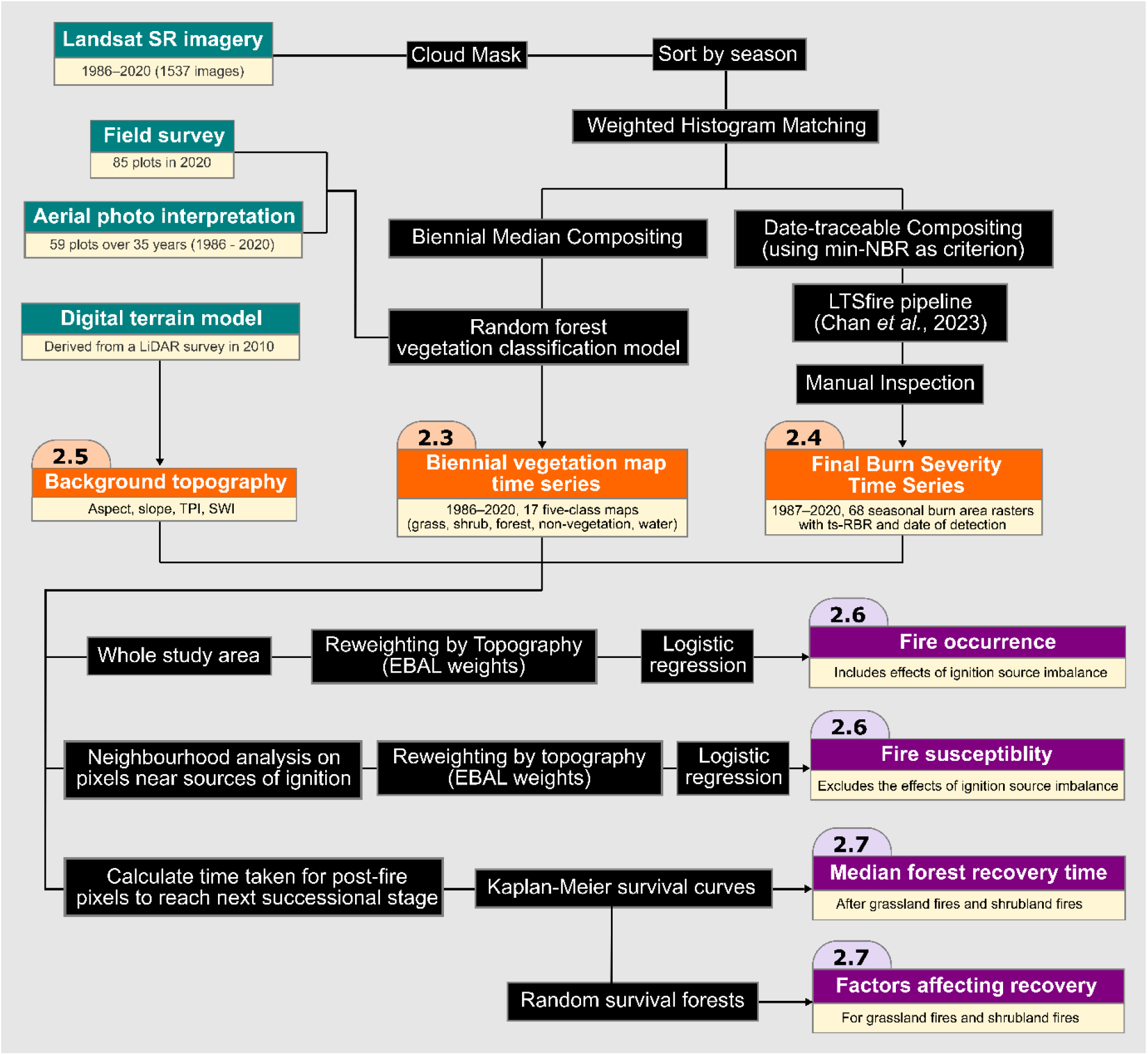
Methodology flow chart showing how fire trap processes were evaluated by remote sensing data. Numbers indicate the relevant section in the main text. Green boxes are input data; black boxes represent steps taken; orange boxes indicate intermediate products; and purple boxes are the fire trap processes quantified.

### 2.3 Vegetation map time series

A Landsat-based vegetation map time series was produced to track changes in vegetation structure over the 35-year study period (1986-2020). We distilled 1537 relevant Landsat surface reflectance scenes into 17 biennial (every two years) composites. Weighted histogram matching, band normalisation and vegetation indices were adopted to make the scenes and composites intercomparable (Wu, 2004; Chan *et al*., 2021, 2023). We then built random forest (RF) classification models based on pixels with known vegetation cover. For every biennial composite, the RF model classified the landscape into five classes (forest, shrubland, grassland, non-vegetation, and water).

Detailed methodology can be found in the **Supporting Information**, with RF model accuracies listed in **Table S1**. Finally, we hypothesised that distance to nearest forest patch may affect post-fire recovery trajectories. Hence, we calculated this distance for all grassland and shrubland pixels using the *distance* function in the *raster* package. Unless otherwise specified, all geospatial analyses were carried out in *R-4.1.0* (R core team, 2021).

### 2.4 Burn area and burn severity time series

Burn areas (BAs) in Hong Kong were mapped across a 35-year Landsat multispectral time series (1986-2020) using the LTSfire pipeline (Chan *et al*., 2023). The pipeline was designed to accurately detect small BAs in regions with high cloud cover and rapid burn area revegetation. The product of LTSfire includes two components – (1) a shapefile of BA polygons with dates of detection and (2) rasters containing estimates of burn severity, time series relativized burn ratio (ts-RBR), for all burnt pixels. We refer readers to Chan et al. (2023) for details on the LTSfire pipeline, but an overview of the pipeline and accuracies can be found in the **Supporting Information** and **Table S2**. We further manually inspected the dataset and removed 1179 dubious features that were potential artifacts and modified the shape of 137 polygons. The final dataset contained 5654 burnt patches.

### 2.5 LiDAR background topography

To investigate how background topography affected vegetation fire susceptibility and post-fire recovery, we built rasters for four topographical variables – slope, aspect, topographic position index (TPI), and SAGA wetness index (SWI). The rasters were generated from a digital terrain model (DTM) based on an airborne LiDAR dataset collected in 2010. Slope and aspect were calculated using the *terrain* function in the *raster* package; TPI was generated by the *gdaldem* function in *GDAL*; while SWI was calculated by calling the *rsaga.wetness.index* function through the *RSAGA* package in *R-4.1.0*. Since many of these topographical variables were resolution-dependent, we selectively downsampled the DTM before generating the topographical layers. Aspect was calculated from a DTM downsampled to 30 m resolution. We then tested how different DTM resolutions affected the TPI and SWI to ensure that the indices (1) did not focus exclusively on highly local topographical features such as boulders or rocks, while (2) not smoothing out the effects of valleys and ridges in the mountainous terrain (**Figure S1**). From the analysis, DTMs of 20m and 15m ground resolutions were used to generate TPI and SWI, respectively. Since local steepness has been reported to affect fire propagation (Viegas and Viegas, 2004), we used the original 1m DTM when calculating slope. After generation, all topographical variables were eventually tidied to 30m ground resolutions to match that of Landsat images. Finally, as the effect of aspect is cyclical, we linearized the variable by calculating cos_aspect with the local optimum aspect of 5.795 in radians (Stage, 1976). The optimum aspect was identified by analysing forest growth rates as measured by photogrammetry- and lidar-based digital surface models (Hong Kong Observatory, 2023) (**Figure S2**).

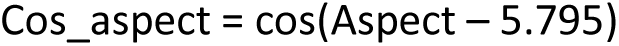

### 2.6 Fire susceptibility, ignition source distribution, and fire occurrence

#### 2.6.1 Overview

Pixel values were extracted from the vegetation, burnt area, and topographical rasters to analyse how different factors shaped the fire regime of the study area. We defined fire susceptibility as the likelihood for pixels to burn given the same exposure to ignition sources. It is an inherent property of the site defined by the vegetation type and background topography. This differs from fire occurrence, which represents the actual proportion of burnt pixels after considering the non-random distribution of ignition sources. We conducted two separate analyses to estimate the differences in (1) fire occurrence and (2) fire susceptibility amongst different vegetation types. By comparing the two estimates, we quantified the contribution of ignition source imbalance on fire patterns. For instance, if grasslands and forests are equally fire prone, but villagers only set fire to easily accessible grasslands not dense forests, we would see a large difference between grassland and forest fire occurrence, while observing no difference in fire susceptibility. The approach avoids the use of indirect proxies, such as distance to roads, to model ignition source distribution. This makes the analysis more robust and generalisable as the relationship between distance to roads and ignition source density is region-specific and dependent on socio-economic factors. Lastly, we also investigated into how topography interacted with vegetation type in determining fire susceptibility on a site level. All analyses below were carried out in *R-4.1.0* (R core team, 2021).

#### 2.6.2 Fire occurrence

For each of the 17 biennial vegetation maps in the time series, we filtered out grassland, shrubland, and forest pixels, then tracked whether the pixels experienced a fire within the two-year window. We also took note of the TPI, TWI, cosine aspect, and slope of the pixels. The *WeightIt* package was used to assign entropy balancing (EBAL) weights to tackle topographical covariate imbalance amongst the four vegetation types (Greifer, 2019). The reweighting process is akin to matching grasslands, shrubland, and forest pixels with comparable background topography, but without discarding or synthetically creating data. We then used the reweighted data to build a logistic regression model that predicted fire occurrence from vegetation type and background topography. Odds ratios were calculated to estimate how fire occurrence differed between different vegetation types as result of both fire susceptibility and ignition source imbalance.

#### 2.6.3 Fire susceptibility

Neighbourhood analysis was performed to estimate how vegetation type and background topography affected fire susceptibility when pixels were exposed to the same ignition source. To achieve this, we focused on the vicinity of detected burn areas where ignition sources were known to have existed. For each burn area polygon, we first located the centroid using the *st_centroid* function in the *sf* package. We then drew the longest line between the centroid and the edge of the polygon. Using the line as the radius, we created a circle representing the theoretical maximum fire extent (orange circle, **Figure S3**). The circle was modified by removing regions that were cut off by non-vegetated areas such as roads or other fire breaks. Pixels within the modified circle would have been exposed to the ignition source, with the vegetation fire susceptibility determining whether it actually burnt. We recognise that pixel fire susceptibility is affected by variables other than vegetation type, and the exposure of the encircled pixels to the ignition source may still vary (directed acyclic graph, or DAG, in **Figure S4**). We addressed these measured and unmeasured confounding variables using *do-*calculus logic (Pearl, 1995, 2009; Shrier and Platt, 2008; Suttorp *et al*., 2015) (see **Supplemental Information** for details). EBAL weights were assigned to the pixels to ensure that pixels of the four different vegetation classes were comparable with respect to the four topographical covariates and distance to fire centroid (Greifer, 2019; Matschinger, Heider and König, 2020; Markoulidakis *et al*., 2022). Using the weighted data, we built a logistic regression model that predicted fire susceptibility from vegetation type, topographical variables, and distance from burn area centroid. All continuous variables were scaled such that the coefficients reflected the effect sizes and variable importance. A forest plot was generated to evaluate variable importance using odds ratios calculated from the model coefficients. The detailed structure of models can be found in **Table S3**.

### 2.7 Survival analysis on post-fire recovery

Post-fire vegetation recovery rates were quantified by running survival analysis on the times it took for burnt pixels to reach the next successional stage. We chose survival analysis as the data is temporal and right censored (Muenchow, 1986; Tepley *et al*., 2018; Therneau, 2019). Two types of right censorship were observed – pixels might have not reached the next successional stage by 2020 or might have experienced another fire. The latter type of censorship was problematic as repeated fires would disproportionally censor pixels that failed to recover. This violated the assumption of non-informative censorship in survival analysis (Therneau, 2019). Hence, we focused our study on the recovery trajectory after the last observed fire between 1986 and 2020. Another assumption made was the unidirectional vegetative succession without retrogressions. The assumption was largely met, with 97.1% of the pixels either staying in the same vegetation class or transitioning to a later successional stage over time (grasslands to shrublands to forests), so we proceeded after filtering out retrogressed pixels.

Median post-fire recovery times after grassland and shrubland fires were estimated by constructing Kaplan-Meier survival curves using the *survival* package (Therneau, 2019). The curves were built from: (1) survival time – the number of years the pixel “survived” as grassland or shrubland before transitioning into forest and (2) censorship – binary variable that records whether a pixel ever became forest in the observed period. The approach worked well for shrubland fires, but for grasslands, it was complicated by the median post-fire recovery time being longer than the 34-year study period. This was problematic as Kaplan-Meier curves are non-parametric and could not be easily extrapolated (Therneau, 2019). To tackle this, we estimated the grassland > young shrubland and young shrubland > forest recovery times separately. “Young shrubland” represented pixels that just transitioned from grassland to shrubland in our vegetation maps. While these pixels have not necessarily experienced a fire in the study period, it is likely that these pixels burnt in the past given the historical fire frequency in Hong Kong (Chan *et al*., 2023). To ensure that the young shrubland pixels have the same topographical profile as the grassland burnt in our study period, we used the *WeightIt* package to generate EBAL weights based on cosine aspect, slope, TPI, and SWI (Greifer, 2019). Finally, we estimated the median grassland to forest recovery time by adding the survival times obtained from the two sets of Kaplan-Meier curves.

To investigate the relative importance of different factors in determining post-fire recovery rates, we used the *randomForestSRC* package to build a random survival forest (RSF) model (Ishwaran, Kogalur and Kogalur, 2023). The RSF model predicted recovery time from pre-fire vegetation type, burn severity (ts-RBR), distance to the nearest forest patch, cosine-aspect, slope, topographical position (TPI), and wetness (SWI). We used RSF as the non-parametric machine learning approach was more robust against multicollinear and non-linear relationships while still being able to handle right-censored temporal data (Ishwaran *et al*., 2008). Variable importance was estimated from the RSF model with subsampling inference and the delete-d jackknife estimator.

To visualise the partial effects of biophysical and topographical variables on post-fire recovery, we constructed 240 Kaplan-Meier curves based on stratified and reweighted datasets. Hypothesising that biophysical and topographical variables may have different effects on grassland and shrubland fires, we analysed grassland to shrubland and shrubland to forest transitions separately. We then stratified the two datasets by ts-RBR, forest distance, TPI, SWI, slope, and aspect, with 20 groups per variable. To isolate the effect of the variable in question, we reweighted the groups to tackle the imbalance of potentially confounding variables. For instance, it is reasonable to expect pixels with low TPI (valleys) being closer to forests and have lower burn severity (ts-RBR), so we assigned EBAL weights to pixels in the 20 TPI groups such that weighted pixels would have the same ts-RBR and forest distance distribution across the groups. Finally, we constructed 20 Kaplan-Meier curves per variable and quantified how changes in the variable affected median post-fire succession time. Further details regarding the Kaplan-Meier curves and EBAL weights can be found in **Table S3**.

## 3. Results

### 3.1 Fire susceptibility, ignition source distribution, and fire occurrence

Our results support the existence of strong fire-vegetation feedbacks (fire traps) in the wet subtropics that were exacerbated by anthropogenic ignition source imbalance. From the neighbourhood analysis, it is estimated that the fire susceptibility of grasslands and shrublands were 19.5 and 8.5 times higher than that of forests, respectively, given the same exposure to ignition sources (**Figure 3**). Ignition sources were not randomly distributed. Grasslands and shrublands were 2.3 and 2.0 times more likely to be exposed to ignition sources compared to forests, respectively. The compounding effects of high fire susceptibility and increased exposure to anthropogenic ignition sources led to much larger differences in actual fire occurrence across vegetation types. Fire occurrence was 44.5 and 16.9 times higher amongst grassland and shrubland pixels, respectively, compared with forest pixels with similar background topographies. Background topography had relatively minor effects on vegetation fire susceptibility. Slope and topographical position had nearly no effect on fire susceptibility (**Figure 3**).

**Figure 3:**
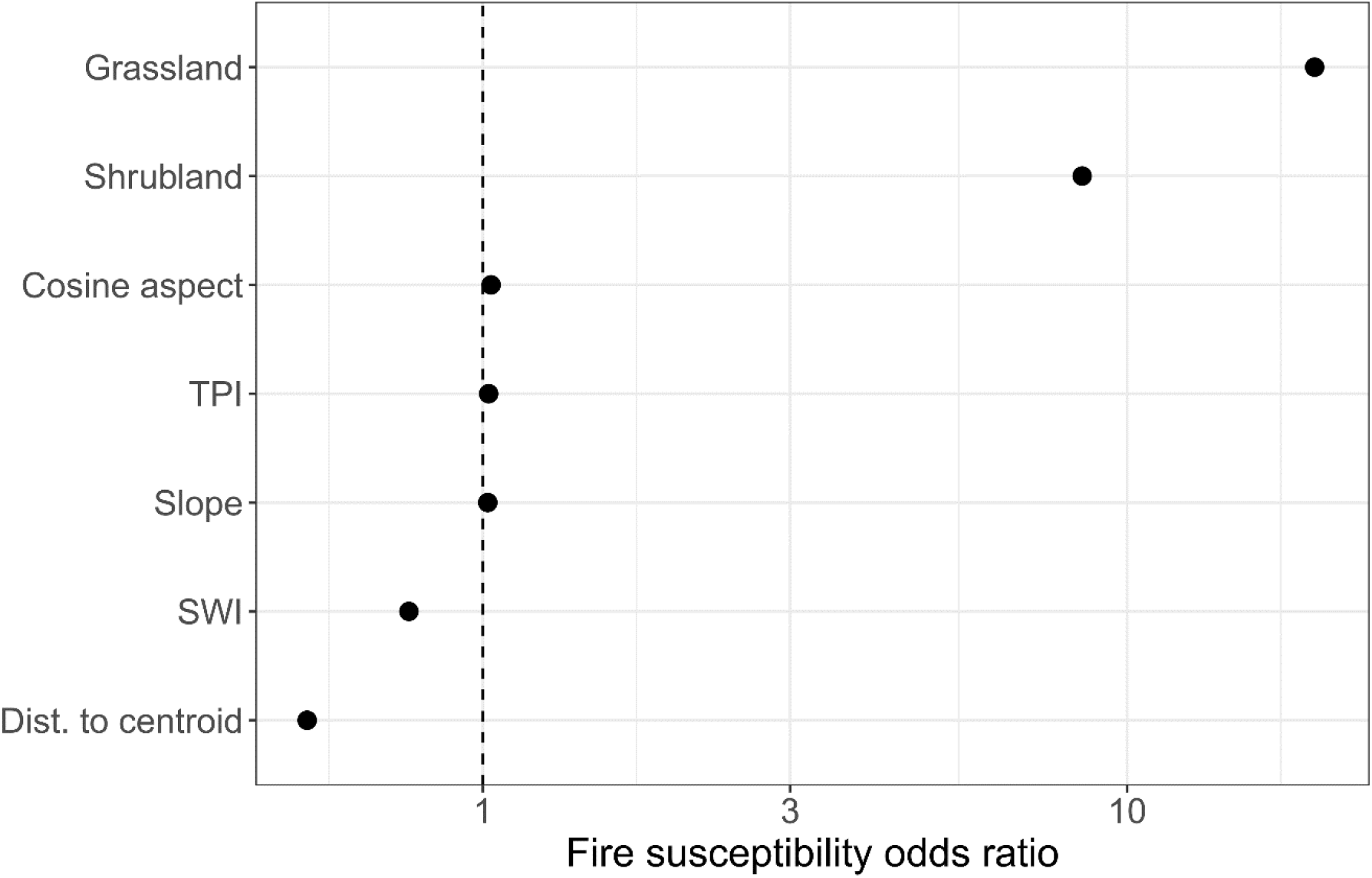
Effects of vegetation type and topographical variables on fire susceptibility in wet subtropical Hong Kong. Odds ratios represent how the variables change the likelihood of a pixel burning, given a fixed exposure to ignition sources. The odds ratios of the two categorical variables (grassland and shrubland) refers to the fire susceptibility compared to that of forests. “TPI” represents topographical position index. “SWI” represents SAGA wetness index. “Dist. to centroid” refers to the distance between the pixel and centroid of the burnt area in the neighbourhood analysis. All pixels in the neighbourhood analysis would have been close to known ignition sources, but a longer distance to centroid would indicate lower exposure at a local level. The 95% confidence intervals were too small to be visible.

Wetter (high SWI) pixels were less fire susceptible (**Figure 4a**) and aspect had vegetation-specific effects on fire susceptibility (**Figure 4b**), but their effect sizes were small compared to that of vegetation type and ignition source abundance (**Figure 3**). Overall, our results show that local fire regimes in the wet subtropics are defined by fire-vegetation feedbacks (fire traps) interacting with ignition source distribution, not background topography.

**Figure 4:**
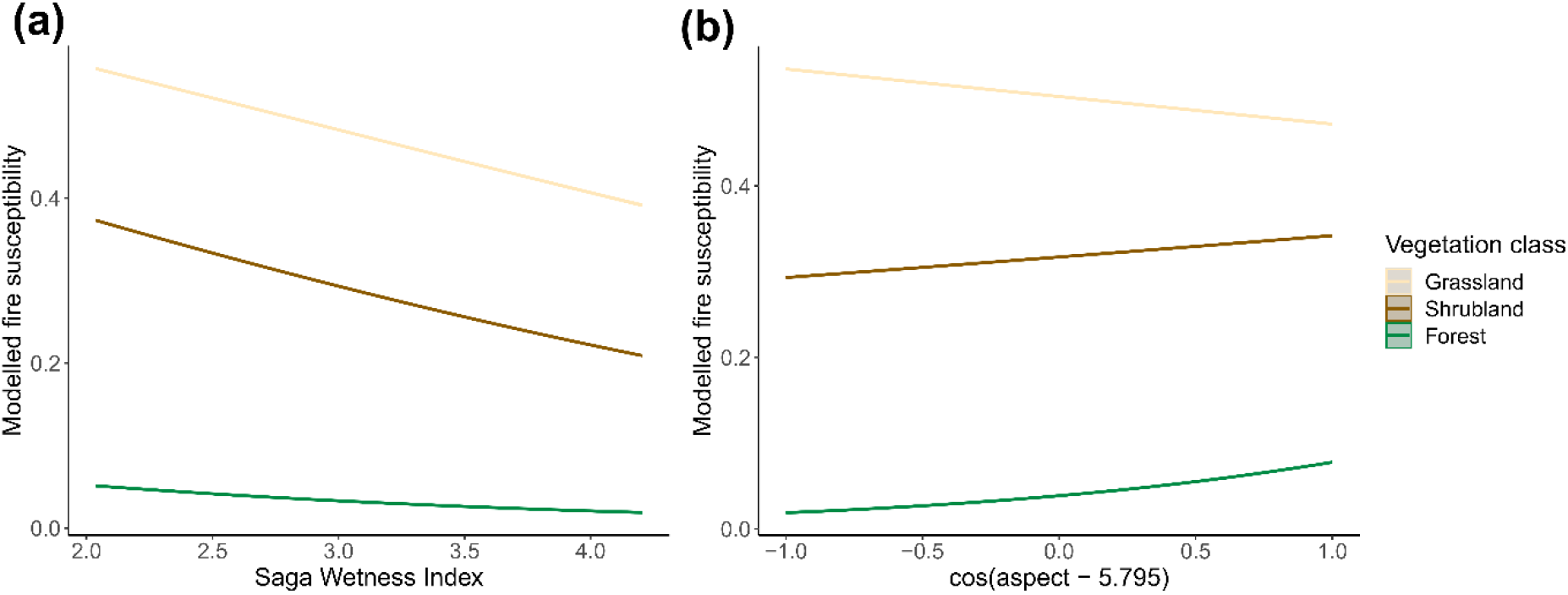
The effects of SAGA wetness index (SWI) and linearised aspect on fire susceptibility modelled by logistic regression. The shaded area (barely visible due to the large sample size) represents the 95% confidence interval.

### 3.2 Post-fire recovery

Pre-fire vegetation type significantly affected post-fire recovery rates, indicating strong legacy effects. The proportion of pixels in each vegetation class before and after detected fires were tallied in **Figure 5**, which shows how pixels moved through the stages of succession over time. Note that while some shrubland pixels retrogressed to grasslands after fires, most tended to stay in the same class. The median time required for grasslands to recover back to forests after fires was 40 years, while shrublands on similar topographies recovered in 19.2 years.

**Figure 5:**
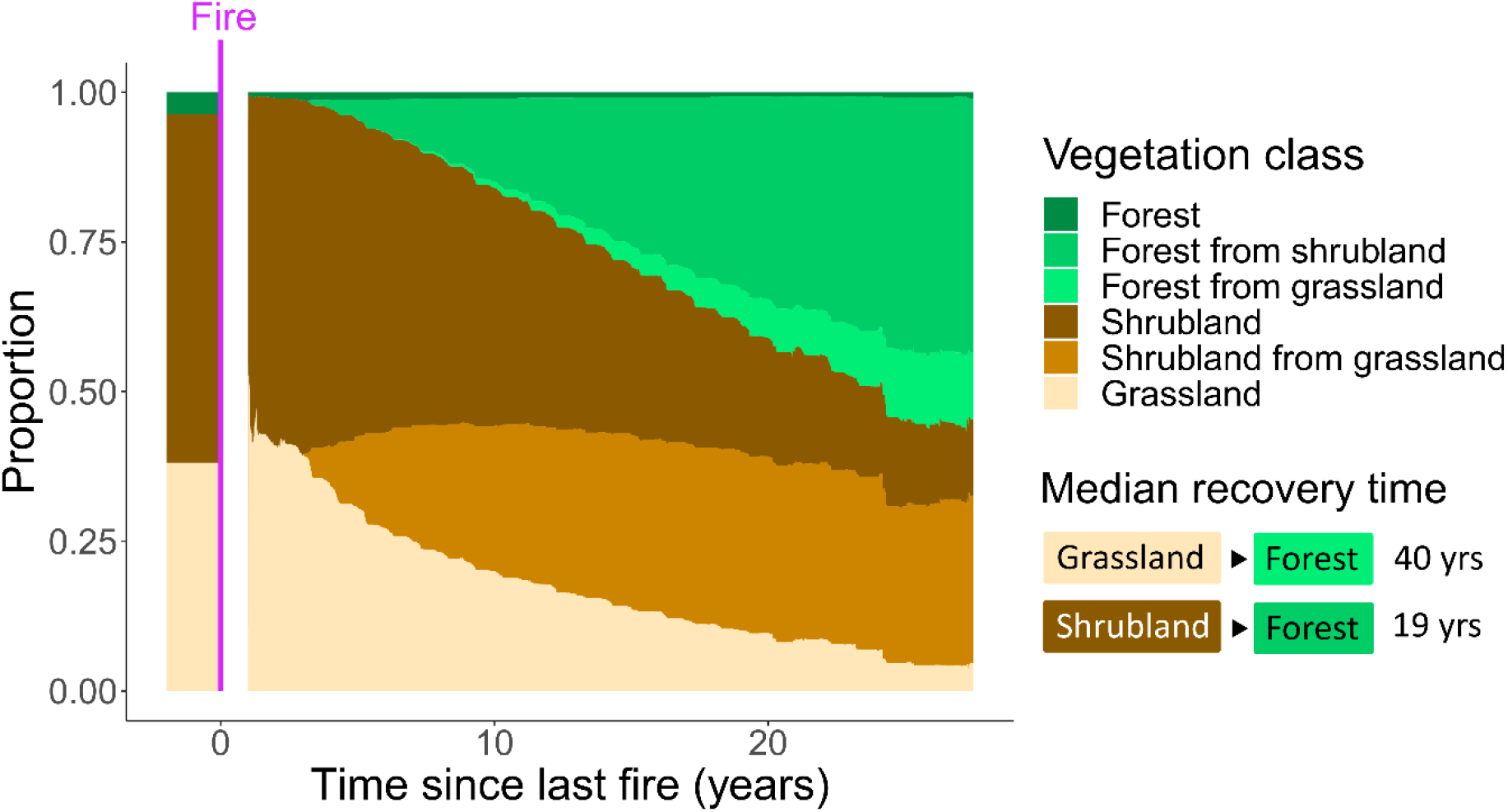
Area plot to visualise post-fire recovery trajectories in wet subtropical Hong Kong. The y-axis represents the proportion of pixels in each vegetation class before (left of the magenta line) and after (right of the magenta line) the fire. The median recovery times were estimated from survival analysis.

Post-fire recovery rates were highly variable and correlated with a range of vegetative, biophysical, and topographical factors (**Figure 6**). Post-fire distance to the nearest forest patch strongly affected rate of recovery (**Figure 6** **and** **Figure 7**). Sites closer to forest patches recovered significantly faster than those further away from forest patches (**Figure S5a/S5b and** **Figure 7**). The effects were strongest for pixels between 0-250 m from forest patch, but levelled off after 250 m. Aspect had a significant cyclical effect on post-fire recovery rates (**Figure 7**). For both grassland to shrubland and shrubland to forest transitions, sites facing the northwest recovered the quickest (**Figure 7**). Interestingly, burn severity was found to have significant but opposite effects on post-fire recovery rates in grasslands and shrublands. Severe fires (high ts-RBR) promoted recovery to shrubland after grassland fires, but inhibited recovery to forests after shrubland fires (**Figure 7**). These effects persisted through the Kaplan-Meier curves (**Figure S5c/S5d**). Finally, pixels in valleys (low TPI) recovered quicker than those on ridges (high TPI), and, given the same topographical position, wetter pixels (high SWI) recovered quicker than drier ones (low SWI). The variable importances of both TPI and SWI were smaller than other variables assessed (**Figure 6**), but both factors seem to be proportionally more important for the transition to shrublands after grassland fires (**Figure 7**). Slope was found to be a non-significant predictor in the RSF model (**Figure 6**), with the slope-stratified dataset generating broadly negative but messy relationships between slope and median recovery times (**Figure 7**). The multitude of factors influencing recovery contrast with the analyses of fire susceptibility, in which a single factor – vegetation type – stood out as the predominant driver.

**Figure 6:**
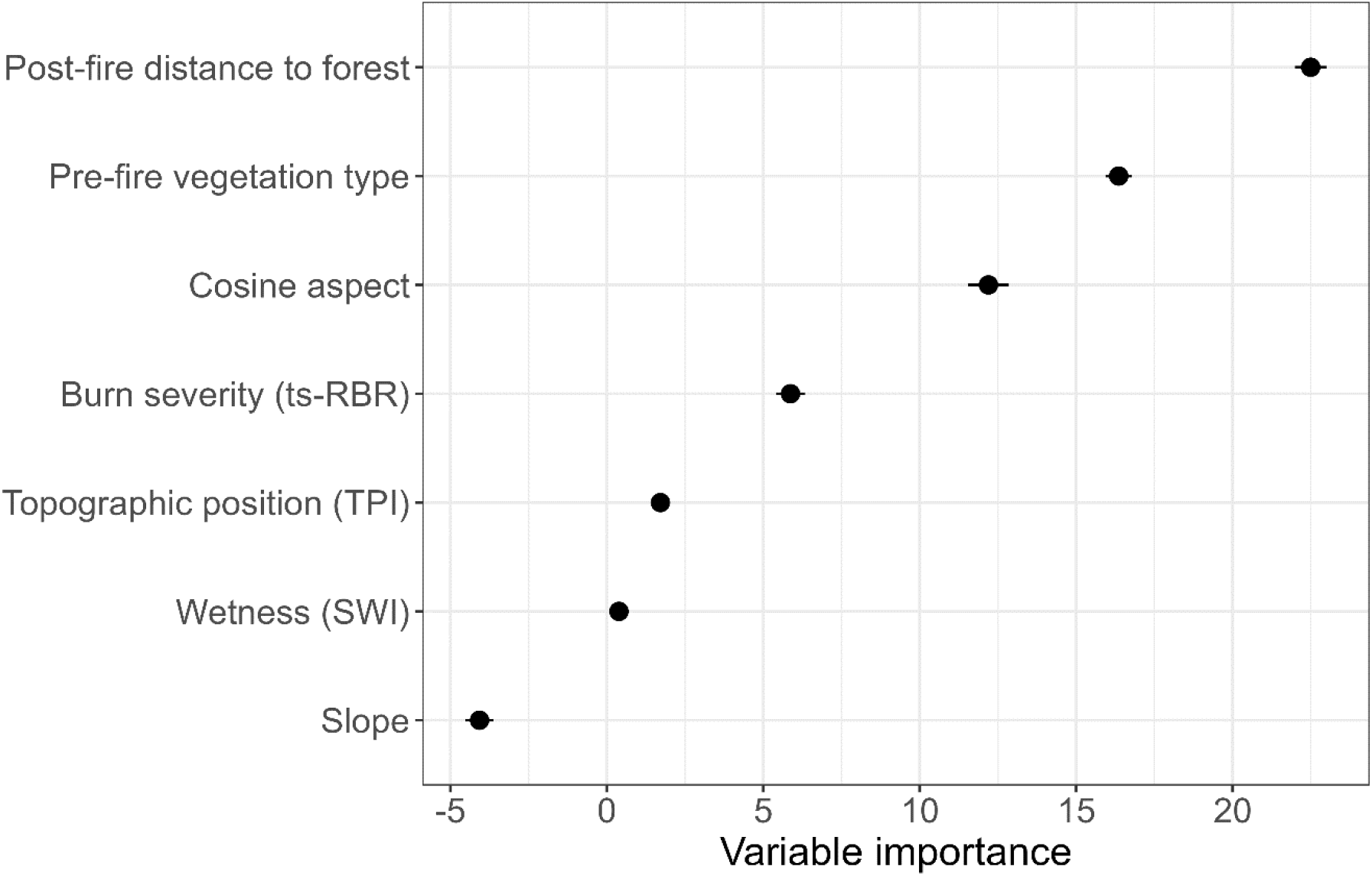
Variable importance derived from a random survival forest (RSF) model that predicts median post-fire recovery times back to forests. All variables were statistically significant (p < 0.05) except for slope. Confidence intervals for variable importances were calculated by subsampling the dataset and using the delete-d jackknife estimator.

**Figure 7:**
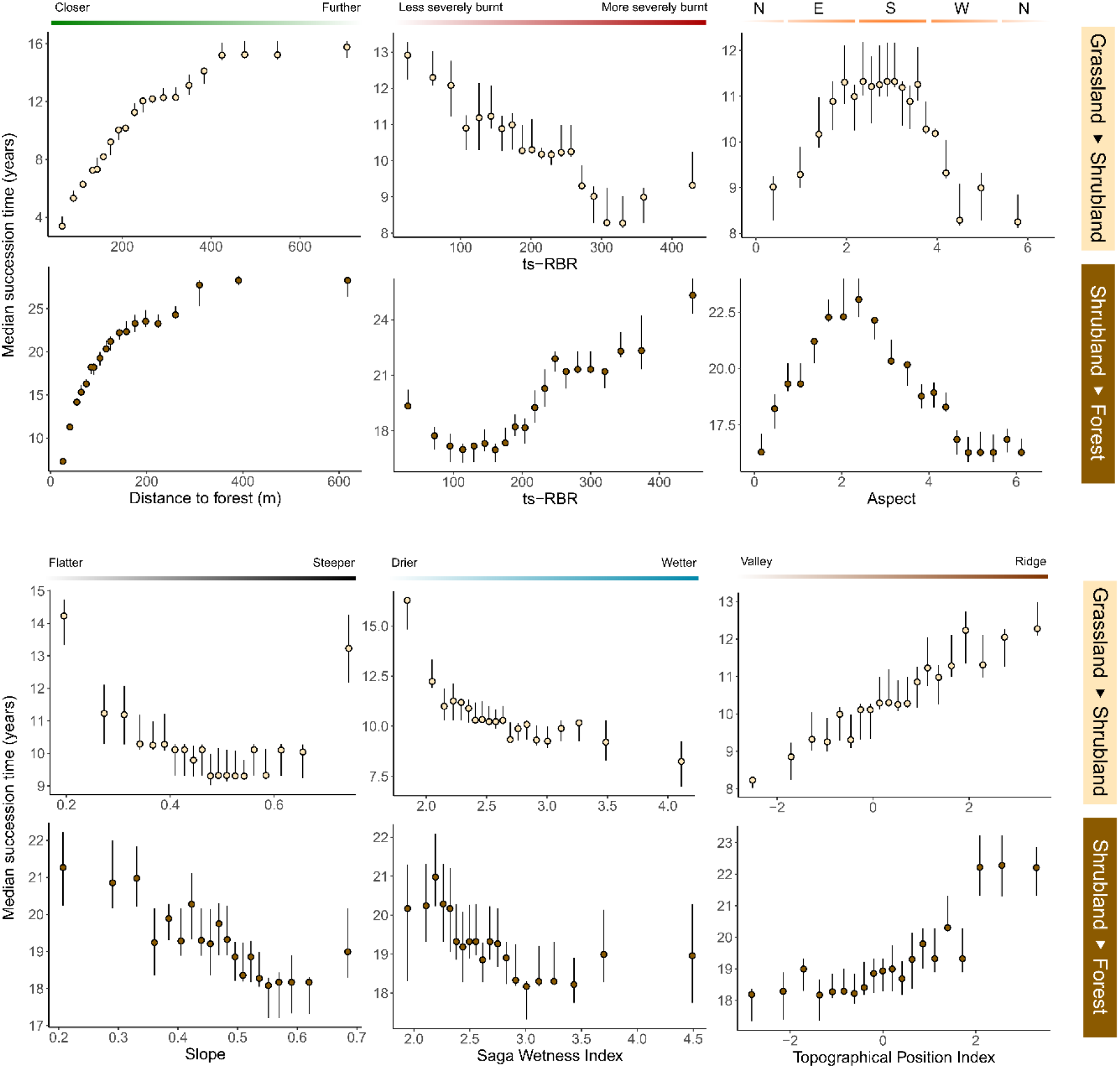
Plots showing how different variables affect median recovery times for the transition from burnt grassland to shrubland and burnt shrubland to forest. The median recovery times were estimated by stratifying the dataset by the variable in question and building Kaplan-Meier curves for each stratified group. Pixels in the groups were assigned entropy balancing (EBAL) weights to tackle the imbalance of relevant covariates to isolate the effect of the variable in question (see Table S2). The error bars represent 95% confidence intervals.

## 4. Discussion

### Quantifying the strength of the fire trap

This study demonstrates the existence of strong fire-vegetation feedbacks in the wet subtropics. Degraded grasslands and shrublands were 20 and 9 times more fire-susceptible than forests. Previous research showed that grassy fuels have three times lower bulk densities compared to litter fuels (Prior *et al*., 2017), and shrubs produce finer litter fuels than forest trees (Plucinski *et al*., 2010). Given that wet subtropical vegetation is not generally fuel limiting, the lower fuel bulk densities in grasslands and shrublands lead to high ignitability and rapid rate of fire spread (Uhl, Kauffman and Cummings, 1988; Hoffmann, Jaconis, *et al*., 2012; Prior *et al*., 2017; Iván *et al*., 2023). Additionally, as more open habitats, grasslands and shrublands tend to retain moisture poorly (Hoffmann, Jaconis, *et al*., 2012; Iván *et al*., 2023). In particular, *Dicranopteris* fern mats and C4 grasses are commonly found in open habitats in Hong Kong. These vegetation types accumulate dead biomass that decompose slowly and desiccate easily, making them particularly fire-prone (Matos, Santos and Chevalier, 2002; Hoffmann, Jaconis, *et al*., 2012). Th high fire susceptibility of early successional vegetation creates fire traps that make degraded wet tropical and subtropical landscapes inherently difficult to restore.

Anthropogenic ignition source imbalance – the tendency for humans to set fire to open grassland and shrubland habitats – greatly exacerbated natural fire traps. In our study area, humans introduced 2.3 and 2 times more ignition sources to grasslands and shrublands, leading to the actual fire occurrences of the two early successional vegetation types being 45 and 17 times higher than in forest patches of comparable background topography. Using distance to roads and settlements as proxies, previous studies also reported the importance of anthropogenic ignition sources in affecting fire occurrence (Oliveira *et al*., 2012; Tien Bui *et al*., 2016). Our results further demonstrates that these ignition sources are not evenly distributed across different vegetation types. Grasslands and shrublands receive more sources of ignition either because they are more accessible to humans or, alternatively, sites closer to settlements tend to be more degraded and less forested.

Surprisingly, background topography hardly affected fire susceptibility. The effect size of the strongest topographical predictor, wetness (SWI), was an order of magnitude smaller than vegetation type and ignition source exposure. This echoes other studies in the region (Tien Bui *et al*., 2016) but is in stark contrast with results from Mediterranean Europe, where several studies reported clear effects of slope or aspect on fire susceptibility, with effect sizes in the same order of magnitude as that of background vegetation type (Carmo *et al*., 2011; Oliveira *et al*., 2013; Barros and Pereira, 2014). These differences highlight how fire regimes in the wet tropics and subtropics fundamentally differ from that in other biomes.

### Factors influencing post-fire recovery rates

Pre-fire vegetation type was the strongest predictor of post-fire recovery rate, with shrublands recovering much quicker than grasslands after experiencing a fire (median recovery time 19 years vs 40 years). Many shrubs and pioneer trees tend to survive the fires and resprout, even in the ecoregions where fires are naturally scarce (Chau, 1994; Van Nieuwstadt, Sheil and Kartawinata, 2001; Teixeira *et al*., 2020). The ability to re-establish itself using basal or epicormic regrowth might partly have evolved against rare natural fires, but it is more likely to be a general trait that helps these species recovery from other disturbances posed by large herbivores or windstorms (Chau, 1994; Teixeira *et al*., 2020).

Other than pre-fire vegetation type, the distance from the nearest forest patch after fires had an equally strong effect on post-fire recovery rates. The relationship between forest distance and recovery time was found to be non-linear and flattens out over distances >250m. This resembles the inverse of seed dispersal kernels, which indicates limitations in seed availability (Levine and Murrell, 2003; Herrmann *et al*., 2016; Rogers *et al*., 2019; Flores and Holmgren, 2021). Unlike regions were serotiny is the norm, burnt patches in the wet tropics and subtropics rely heavily on external seed sources to move along the recovery trajectory (Van Nieuwstadt, Sheil and Kartawinata, 2001; Flores and Holmgren, 2021). Au et al. (2006) quantified the seed rain in degraded grasslands and shrublands in the study area and found large variations in the number of seeds per m^2^ per year, ranging from 47 in open grasslands to >6000 under female shrubs on grasslands. While one might argue that 47 seeds per m^2^ per year might be sufficient to push the landscape through succession, many seeds would fail to penetrate the dense mats of early successional grasses or ferns to reach the mineral soils, and established seedlings can get smothered under the thick vegetation (Pang *et al*., 2018; Rochimi, Aj and Meador, 2021). Many seeds may also belong to shorter shrub species that may not necessarily help the site recover back to forests. Most species in the four dominant tree families in Hong Kong (Lauraceae, Moraceae, Fagaceae, Euphorbiaceae) rely on animals for seed dispersal (Dudgeon and Corlett, 2004). The lack of perch sites for birds and suitable habitats for scatter-hoarding mammals could prevent the seed dispersal kernels from extending into large burnt areas far from forest patches (Levine and Murrell, 2003; Au, Corlett and Hau, 2006; Rogers *et al*., 2019).

Interestingly, high burn severities promoted post-fire recovery in burnt grasslands but inhibited recovery in burnt shrublands (**Figure S5c, S5d, and** **Figure 7**). Past studies have generally found post-fire recovery times to be longer for more severely burnt sites, even if recovery rates were higher (Ireland and Petropoulos, 2015; Bartels *et al*., 2016; Bright *et al*., 2019). Severe fires tend to cause higher plant mortality, especially for fire-sensitive late successional tree saplings (Hoffmann, Schroeder and Jackson, 2003; Bright *et al*., 2019). This corroborates with the patterns observed in shrublands in our study (**Figure S5d and** **Figure 7**). The opposite relationship observed in grasslands was, however, unexpected. One possible explanation is that dense grasslands might have arrested succession (Rochimi, Aj and Meador, 2021). More severe fires could open the habitat for shrub or tree encroachment, though targeted field surveys would be needed to test this hypothesis.

Background topography also had strong influences on post-fire recovery rates. Echoing previous studies conducted in the northern hemisphere (Pausas and Vallejo, 1999; Wittenberg *et al*., 2007; Ireland and Petropoulos, 2015), we found post-fire recovery to be faster on north-facing slopes. This could be attributable to more sheltering on north-facing slopes, which dampen local fluctuations in temperature and humidity (Stage, 1976; Stage and Salas, 2007; Ireland and Petropoulos, 2015). We would also expect such sheltering to benefit east-facing slopes, as it warms up quicker in the morning but avoids overheating during the day (Stage, 1976). The optimal aspect for post-fire recovery was, however, skewed to the northwest in Hong Kong (**Figure 7**). This might be due to the prevailing wind direction in Hong Kong from the southeast, which leaves westward slopes better sheltered (Hong Kong Observatory, 2023). We also found that sites in valleys (low TPI) and wetter areas (high SWI) had quicker post-fire recovery. Research have demonstrated that DTM-based position and wetness indices serve as effective proxies of variations of microclimates (Jucker *et al*., 2018; Man *et al*., 2022; Marsh *et al*., 2022; Marsh, Krofcheck and Hurteau, 2022). A recent study by Marsh, Crockett, et al. (2022) found DTM-based topographical variables to be almost as accurate as direct microclimatic measurements in predicting post-fire seedling survival in New Mexico. The study reported high seedling survival in areas with high topographical wetness and low TPI, which corroborates with patterns in post-fire recovery observed in Hong Kong despite the large climatic differences between the two study systems. Our results further suggest that TPI and SWI may be affecting post-fire recovery in slightly different ways despite the two variables being correlated with each other. TPI did not fully capture the variation in wetness. Even after reweighting by TPI, SWI was still negatively correlated with recovery times (**Figure 7**). SWI may have captured edaphic factors better (e.g. soils in slopes at the foot of mountains may be wetter than valleys near mountaintops) (Böhner and Selige, 2006). On the other hand, TPI was a more important variable for predicting recovery times in the RSF model (**Figure 6**), indicating that it might have captured information other than wetness, such as sheltering from wind or direct sunlight (Dobrowski, 2011; Jucker *et al*., 2018). Interestingly, wetness seems to have a stronger effect on the grassland to shrubland transition than in the shrubland to forest transition (**Figure 7**). Research in nearby Guangdong suggests that shrubs act as both facilitators and competitors to tree saplings in wet subtropical environments (Liu *et al*., 2013). In drier sites, shrubs moderate post-disturbance microclimates by reducing irradiance and ameliorate temperature fluctuations (Liu *et al*., 2013; Urza *et al*., 2019; Crockett and Hurteau, 2022). In wetter sites, shrubs compete with tree saplings and undermine the topographical benefits (Liu *et al*., 2013). Our results indicate that these biotic buffering effects of shrubs may have overridden topographical determinants of post-fire recovery rates, which, over a landscape scale, might smooth out spatial variations in rates of forest establishment after shrubland fires.

### Escaping the fire trap

Fire traps tend to maintain early successional vegetation once wet subtropical landscapes are degraded, but several policy actions could help overcome these traps. While our results supported the existence of strong natural fire-vegetation feedbacks, it also revealed the observed difference in fire occurrence to be partly anthropogenic due to ignition source imbalance. In other words, fire suppression campaigns would not only reduce fire occurrence across all habitats but would also disproportionally reduce fire occurrence in early successional vegetation. Hong Kong also provides a valuable case study for how fire suppression should be carried out in the wet tropics and subtropics. Over the past 70 years, the government set up a Fire Danger Warning System based on weather and fuel conditions (Hong Kong Observatory, 2023). Public education campaigns were launched to prompt citizens to properly handle potential ignition sources and heed any fire warnings. The government then established a network of 11 fire lookouts on mountaintops such that fires are quickly detected and tackled. The results of these efforts were substantial, with yearly fire occurrence more than halved over the last three decades (Chan *et al*., 2023). While the approach would be costly to implement over larger areas in developing countries, it nevertheless provides a viable pathway towards effective fire-suppression. Fires could also be managed by controlling their spread. Fire breaks can stop ignition sources in other areas from spilling over to sites designated for restoration (Scheper, Verweij and van Kuijk, 2021). A common approach to establish fire breaks is to remove vegetation across a strip of land to create “fuel breaks” (Shinneman *et al*., 2019). Our results, however, suggest that this might be ineffective in the wet subtropics. Vegetative regrowth on fuel breaks is fast in wet ecoregions, which makes them difficult to maintain (Rochimi, Aj and Meador, 2021; Scheper, Verweij and van Kuijk, 2021; Chan *et al*., 2023). If not properly maintained, grasses and shrubs would be highly susceptible to fires regardless of the background topography of the fuel break (**Figure 4**). Green fire breaks may provide a viable alternative in the wet tropics. These fire breaks are created by planting strips of secondary forest or connecting fragmented forest patches (Curran *et al*., 2017). This could be effective in the wet subtropics by levying the fire-vegetation feedback and the fire-resistance of close-canopied forests to stop fire spread, though the ideal widths of these breaks would need to be determined by further investigation and experimentation. Lastly, since fire susceptibility drops sharply across the stages of vegetation succession, fire suppression would become easier and less costly over time. To help landscapes escape fire traps, resources should be disproportionally allocated to suppress fires at the start of restoration projects.

Results from this study also have direct implications on vegetation management after fires. Firstly, rather than using the rebound time of vegetation indices such as NDVI, we described post-fire recovery by estimating recovery time back to different stages of succession. This is much more relevant for land managers hoping to restore the landscape past its pre-fire degraded condition. Secondly, unlike fire-susceptibility, post-fire recovery in the wet subtropics is significantly affected by a range of vegetation, biophysical, and topographical factors. Land managers may consider using the results to perform direct seeding in regions where natural seed sources are rare. Resources could be redirected to replant burnt sites that would not have naturally recovered within a reasonable time frame (Rurangwa *et al*., 2021; Law *et al*., 2023). Alternatively, under budget limitations, managers could also consider using the model to identify areas that could readily undergo natural regeneration. These areas could be prioritised and protected from further disturbances before attempting to restore areas that require active interventions.

## 5. Author contributions

Both authors conceived the ideas and designed methodology; Aland H. Y. Chan collected the data; Aland H. Y. Chan analysed the data; Aland H. Y. Chan led the writing of the manuscript. Both authors contributed critically to the drafts and gave final approval for publication.

## Acknowledgements

This work was supported, in whole or in part, by the Gates Cambridge Trust (OPP1144). Under the grant conditions of the Trust, a Creative Commons Attribution 4.0 Generic License has already been assigned to the Author Accepted Manuscript version that might arise from this submission. We would also like to thank the Civil Engineering and Development Department, the Fire Services Department, and the Agricultural, Fisheries and Conservation Department for providing LiDAR and fire data used in the study.

## 7. Data availability statement

Data made publicly available in Figshare (https://doi.org/10.6084/m9.figshare.c.6668111).

