## Supporting Information for "Fire traps in the wet subtropics: new perspectives from Hong Kong"

**Methods to create the Landsat-based vegetation map time series**

The five-class vegetation map time-series underpinning the study was created from a total of 1537 Landsat 5, 7, and 8 surface reflectance (SR) scenes captured between 1986 and 2020. The scenes were downloaded through Google Earth Engine (Gorelick *et al.*, 2017). Clouds were masked out using the in-built cloud bitmask and by removing pixels with mean RGB reflectances >0.2. We carried out weighted histogram matching to make the scenes more intercomparable to one another (Chan *et al.*, 2023). The scenes were then distilled into 17 biennial (every two years) median composites. We created separate composites for winter (November – February) and summer (March – October) as winter browning is locally important for distinguishing grasslands from other vegetation types. To minimise the effects of pixel brightness on classification accuracy, two vegetation indices (VIs), the normalised difference vegetation index (NDVI) and enhanced vegetation index (EVI), were added as extra bands to the raster. Additionally, the reflectances of the six Landsat bands were normalised by dividing the reflectance of each band by the mean reflectance of all the bands for any given pixel (Wu, 2004; Chan *et al.*, 2021).

A training dataset was created by extracting the reflectances and VIs from sites with known vegetation and landcover types. The vegetation and land cover of these sites were determined by a combination of field data and historical aerial photo interpretation. Field data consisted of 85 plots, each 10x10 m in size, selected from a vegetation survey conducted in 2020. The plots were grouped into forest, shrubland, or grassland classes based on species composition. A canopy height model (CHM) derived from LiDAR data collected by the Civil Engineering and Development Department (CEDD) was used to ensure that all forest plots had median heights >3m, following Abbas et al. (2016). We are aware that some studies adopt a higher 5m threshold for forests (Di Gregorio, 2005), but forests in Hong Kong tends to be of lower statue due to frequent typhoons, and forests would have grown significantly between the LiDAR survey in 2010 and vegetation survey in 2020. An additional 59 points were collected by interpretating historical aerial imagery. The images used were collected by the Lands Department to periodically cover the entire territory of Hong Kong over a 59-year period (1963–2021). Accessible images were mostly not orthorectified, but the high spatial resolution (up to 10 cm) made it possible to locate patches of grasslands, shrublands, or forest by matching local geological features. The final training dataset consisted of data extracted from 144 (85 field plots + 59 historical points) pixels.

Based on the training dataset, we then built a supervised random forest (RF) model that classified pixels into the five vegetation and landcover classes, namely forest, shrubland, grassland, water, and non-vegetation. The accuracy of the model was assessed by 17-fold cross validation. Validation was carried out across years (i.e. training and validation data were never from the same set of Landsat composites). The results of the cross-validation exercise could be found in **Table S1**. Finally, we built an RF model based on the entire training dataset. This model was applied to all 17 biennial LS composite to produce the five-class vegetation map time series (1986-2020).

**A note on Plantations**

Plantations were not analysed as a separate vegetation class in this study. Plantations in Hong Kong are not commercial and were mainly established in Hong Kong to accelerate restoration of degraded landscapes. Many older plantations consist of monocultures of *Lophostemon confertus*, *Acacia confusa*, and *Pinus elliottii*. In recent years, however, native trees are increasingly used to create mixed-species forests. We recognise that plantations could have different fire properties as native forests. However, these properties are dependent on the species composition, and relevant records do not currently exist. In practice, plantations are difficult to distinguish from native forests or shrublands based on Landsat imagery (Kwong *et al.*, 2022). Since plantations only accounts for a small proportion of the landscape (Kwong *et al.*, 2022), we decided against analysing them separately.

**Accuracy of the LTSfire product**

LTSfire is a burn area (BA) detection algorithm designed to map BAs in challenging wet tropical and subtropical regions with high cloud cover and rapid post-fire revegetation (Chan *et al.*, 2023). In the pipeline, we first uniformised the 1537 Landsat 5, 7, 8 surface reflectance scenes by weighted histogram matching (Section 2.2.1). We then produced a total of 68 seasonal composites from the scenes. Minimum normalised burn ratio (min-NBR) was used as the criterion for compositing to highlight burnt areas. The scenes were also composited in a date-traceable manner, with dates of capture carried forward from scenes to composites. Random forest (RF) models were then trained to predict pixel resemblance to BAs based on 94 known BAs. The RF model predictions were later thresholded twice to get seed polygons and growth polygons, which were iteratively merged into a final BA product. Burn severity of pixels encircled by the BA polygons were estimated by the time series relativized burn ratio (ts-RBR), a modified version of the relativized burn ratio (RBR) (Parks, Dillon and Miller, 2014) for time series data.

The accuracy of the LTSfire product was estimated through a 10-fold cross validation with reference to known burn areas (BAs) cataloged in government databases. We specifically ensured that the training and validation dataset consisted of pixels from different fire events, not from different parts of the same BA. We also used these known BAs to evaluate global MODIS- and Landsat-based BA products, including MCD64A1 released by NASA, MCD64A1 produced by ESA, and GABAM by Long et al. (2019). Overall, the omission error of LTSfire was low (11%) compared with global BA products such as GABAM (49%), MCD64A1 (72%), and FireCCI51 (96%). Commission errors were also low, accounting for 0.5-2.4% of total area. Temporally, most estimated fire dates were accurate to within a month of the actual fire, and 76.9% of dates were accurate to ±2 months of the actual fire. Further details on the LTSfire pipeline and product validation could be found in Chan et al. (2023).

**Table S1**: Confusion matrix showing the accuracy of the random forest (RF) vegetation classification model. Accuracies were assessed by 17-fold cross validation of training data from different years. The overall accuracy was 0.87 with a Kappa of 0.84.

| | | Reference | | | | | User's accuracy |
|---|---|---|---|---|---|---|---|
| | | Forests | Grasslands | Non-vegetation | Shrublands | Water | |
| **Prediction** | Forests | 38 | 1 | 0 | 3 | 0 | 0.91 |
| | Grasslands | 0 | 26 | 0 | 1 | 0 | 0.96 |
| | Non-vegetation | 0 | 0 | 26 | 0 | 0 | 1 |
| | Shrublands | 3 | 14 | 0 | 36 | 0 | 0.68 |
| | Water | 0 | 0 | 1 | 0 | 27 | 0.96 |
| **Producer's accuracy** | | 0.93 | 0.63 | 0.96 | 0.9 | 1 | |

**Table S2**: Accuracy of the LTSfire burn area (BA) product compared to that of other BA products (adapted from Chan et al. (2023)). The accuracies were estimated by 10-fold cross validation or direct comparison with manually delineated BA polygons in government fire records. Site omission error refers to the proportion of BA polygons omitted. Area omission error refers to the proportion of omitted BA pixels. Commission error refers to the proportion of unburnt pixels being misclassified as burnt. Numbers in bold represents the highest accuracy or lowest error. More details on the training and validation of the BA datasets can be found in Chan et al. (2023).

| Dataset | Overall Accuracy | Site Omission Error | Area Omission Error | Commission Error |
|---|---|---|---|---|
| LTSfire | **0.952** | **0.0319** | **0.112** | 0.0242 |
| GABAM | 0.860 | 0.565 | 0.493 | 0.012 |
| FireCCI51 | 0.720 | 0.987 | 0.960 | **0** |
| MCD64A1 | 0.799 | 0.949 | 0.720 | **0** |

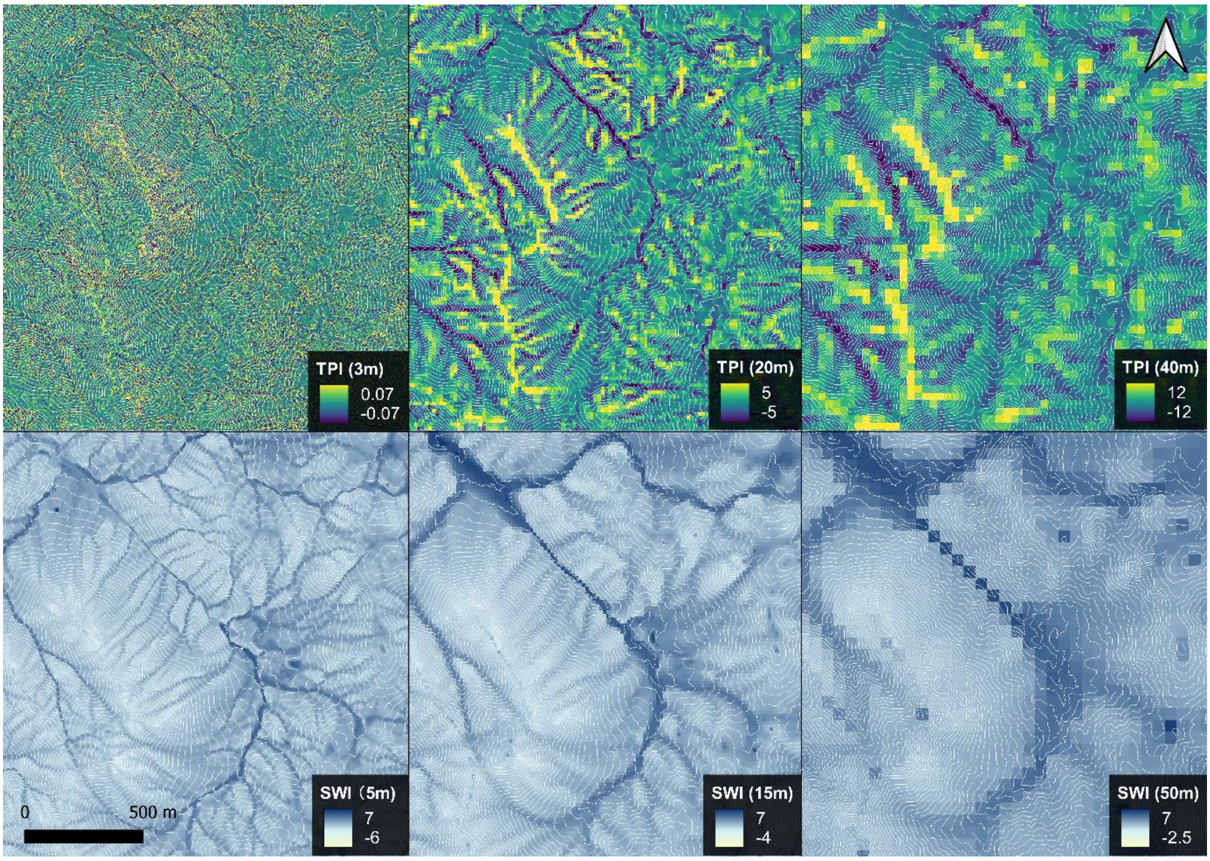


**Figure S1**: The effect of resolution on TPI and SWI.

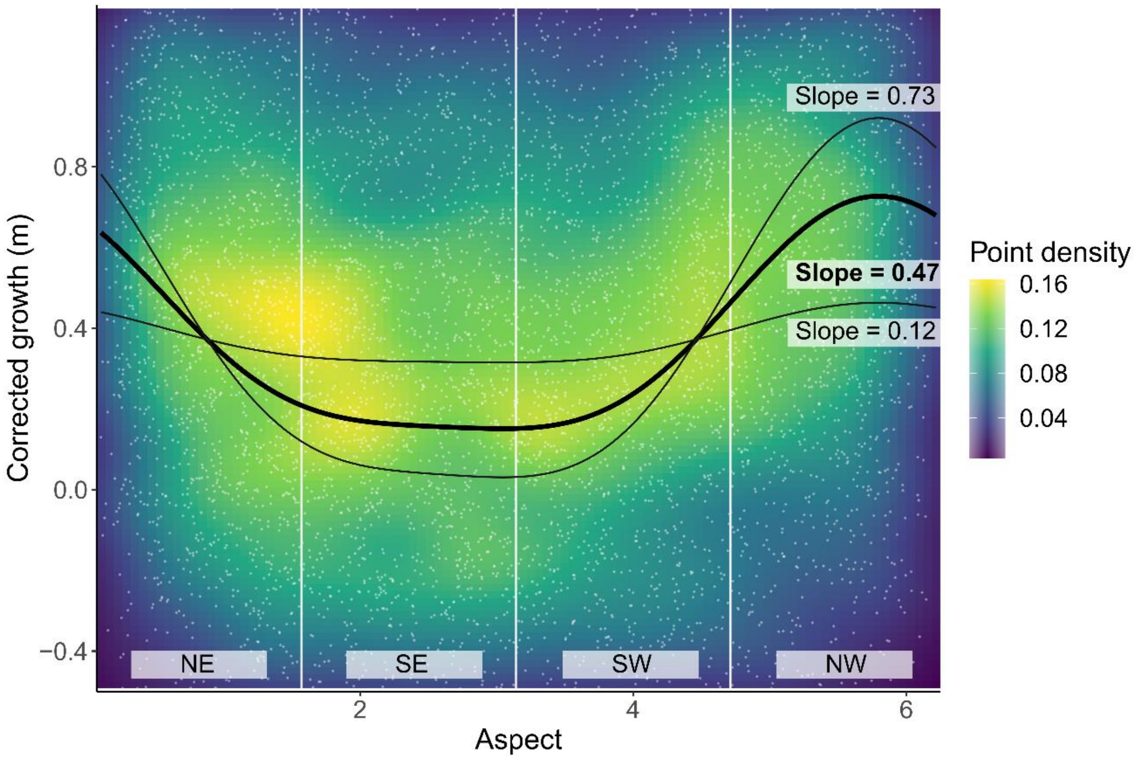


**Figure S2**: The local optimal aspect for vegetative growth in Hong Kong lies in the northwest (5.795 radians). The points represent the change in height of vegetation between two digital surface models in 2010 and 2014. The background colour represents the point density. The black lines represent the mathematical model outlined by Stage (1976) with parameters adjusted to fit the data

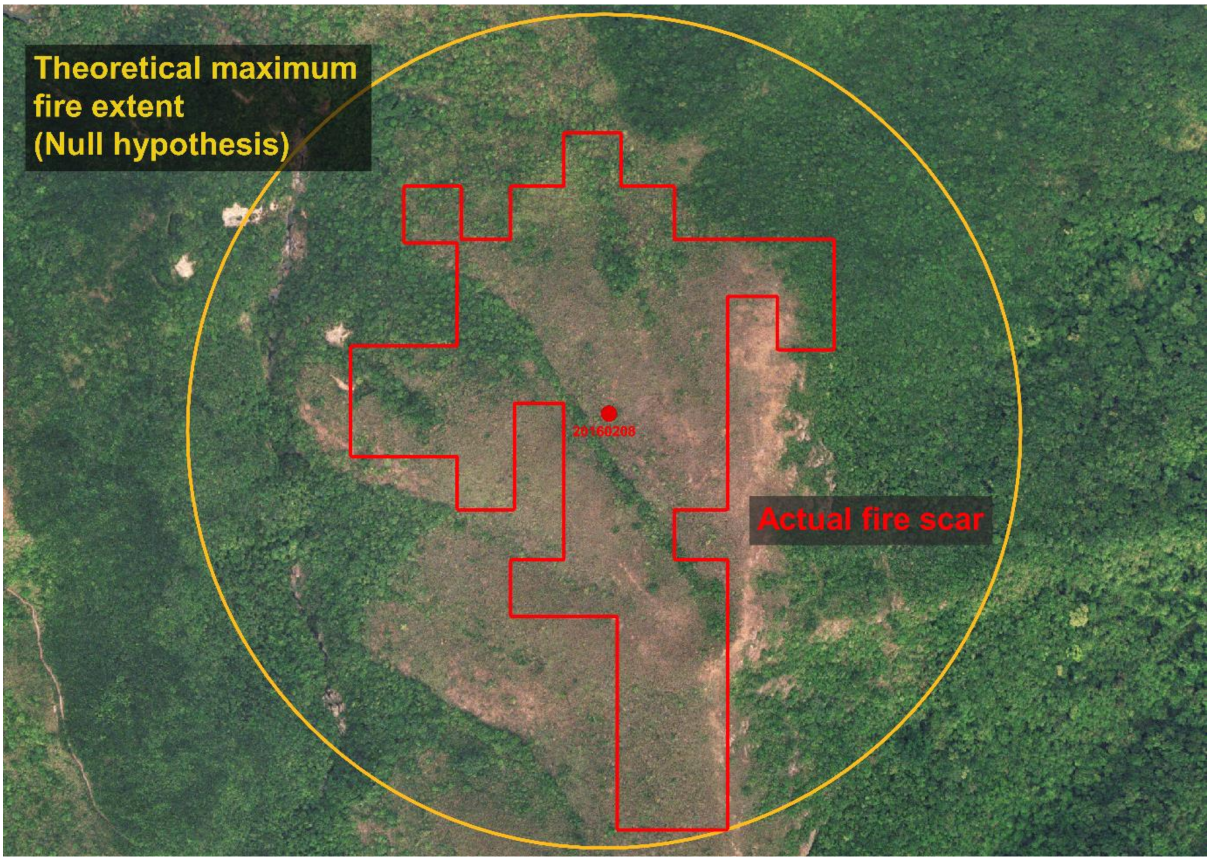


**Figure S3**: Neighbourbood analysis to estimate fire susceptibility.

### A note on covariate imbalance and *do*-calculus

When attempting to estimate the difference in fire susceptibility amongst different vegetation types (grasslands, shrublands, and forests), covariate imbalance could affect the interpretation of the results. For instance, forests could disproportionately occupy wetter valleys, and its low fire susceptibility may be due to topography not vegetation type. Another covariate that remained was the difference in exposure to ignition sources. Although we selected pixels in the neighbourhood of existing burnt areas, which ensured that all pixels have some level of exposure to the source of ignition, the different vegetation types could still have some residual differences in ignition source exposure. It is reasonable to speculate that forests could be further away from the source of ignition amongst all pixels in the circle (**Figure S3**). A fair comparison of fire susceptibility between the different vegetation types could be achieved by reweighting pixels such that the reweighted pixels have the same statistical distribution of covariate values across vegetation types.

When a large number of variables are present, it is often difficult to keep track of which variables need to be addressed or controlled. Pearl (1995) pointed out that it is often unnecessary to control for all variables that could potentially confound our inference. Rather, we could use a graphical approach with a set of logical steps (*do*-calculus) to identify the correct variables to address before testing how the exposure variable (independent variable) affects the response variable (dependant variable). The approach involves first drawing a directed acyclic graph (DAG) that includes the exposure variable, response variable, and all other relevant measured or unmeasured variables. We connect the variables by how they relate to each other. The goal is to identify the paths that connect the exposure variable to the response variable (also known as "back doors"). These paths need to be addressed and blocked by approaches such as matching or reweighting. Each of these paths only has to be blocked once, which would be necessary and sufficient to make reliable causal inference between the two variables of interest (Shrier and Platt, 2008; Pearl, 2009; Suttorp *et al.*, 2015).

**Figure S4** shows the DAG for the study of fire susceptibility amongst different vegetation types and how variables are controlled to minimise the bias when making the causal inference.

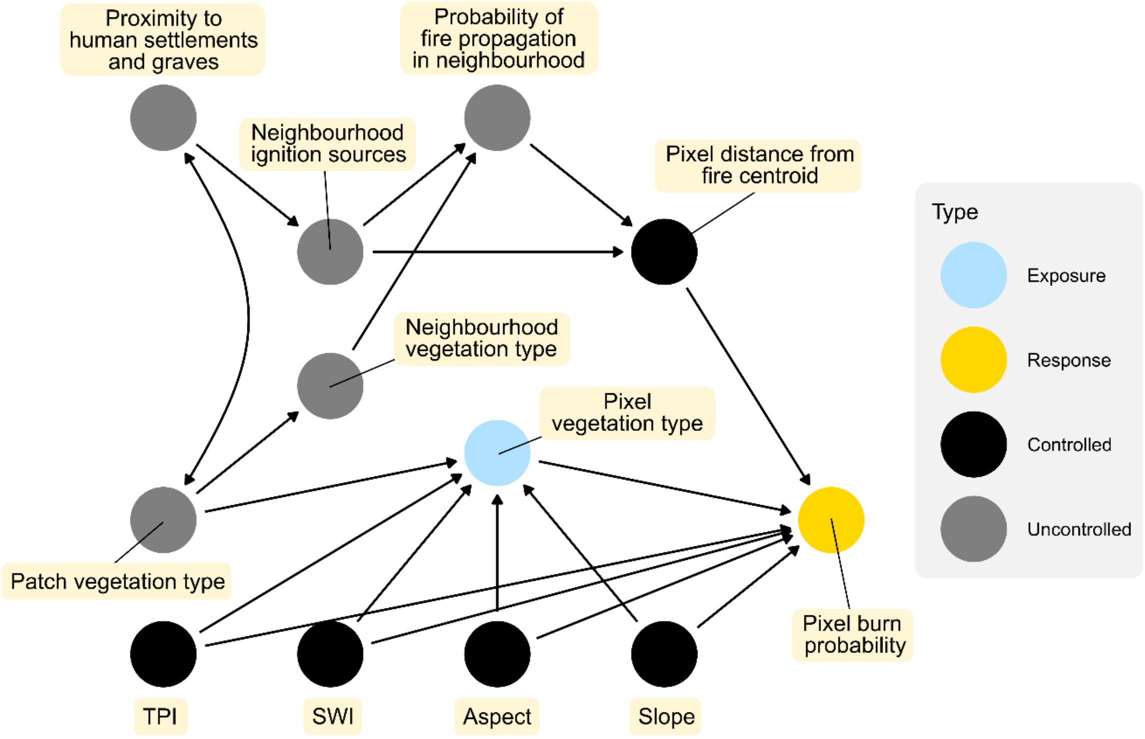


**Figure S4**: Directed acyclic graph (DAG) of the study on fire susceptibility amongst different vegetation types

**Table S3**: Structure of models built. The section number corresponds to where the model was described. TPI refers to topographical position index; SWI refers to SAGA wetness index; cos_aspect refers to aspect linearised by a cosine function; EBAL refers to entropy balancing weights.

| Section | Model | Predictor variables | Outcome variable | Notes |
|---|---|---|---|---|
| 2.5.2 | Logistic regression | Vegetation type (Forest, shrubland, grassland), TPI, SWI, cos_aspect, slope | Probability of the pixel experiencing a fire | For estimating how fire occurrence differs between different vegetation types (no correction for ignition source imbalance). Training pixels was assigned EBAL weights to minimise covariate imbalance between different vegetation types. |

| | | | | |
|---|---|---|---|---|
| 2.5.3 | Logistic regression | Vegetation type (Forest, shrubland, grassland), distance from burn area centroid, TPI, SWI, cos_aspect, slope | Probability of pixel burning given fire source in neighbourhood | For estimating how different variables affect fire susceptibility after correcting ignition source imbalance (**Figure 3**). Training pixels was assigned EBAL weights to minimise covariate imbalance between different vegetation types. Interaction terms were not included here for better visualisation. |
| 2.5.3 | Logistic regression | Vegetation type (Forest, shrubland, grassland), distance from burn area centroid, TPI, SWI, cos_aspect, slope, interaction terms between vegetation type and other variables | Probability of pixel burning given fire source in neighbourhood | For estimating how different variables affect fire susceptibility (**Figure 4**). Training pixels was assigned EBAL weights to minimise covariate imbalance between different vegetation types. |
| 2.6 | Kaplan-Meier survival function | NA | Burnt shrubland recovery rate (years to 50% forest) | For estimating median recovery time after shrubland fires |
| 2.6 | Kaplan-Meier survival function | NA | Burnt grassland recovery rate to young shrubland (years to 50% young shrubland) | For estimating median recovery time after grassland fires |
| 2.6 | Kaplan-Meier survival function | NA | Young shrubland recovery rate (years to 50% forest) | For estimating median recovery time after grassland fires |
| 2.6 | Kaplan-Meier survival function | Stratified TPI | Burnt grassland to young shrubland succession rate<br><br>Burnt shrubland to forest succession rate | Stratified into 20 TPI groups for **Figure 7**. Pixels were assigned EBAL weights such that each TPI stratum had the same distribution of ts-RBR and distance from nearest forest patch |
| 2.6 | Kaplan-Meier survival function | Stratified SWI | Burnt grassland to young shrubland succession rate<br><br>Burnt shrubland to forest succession rate | Stratified into 20 SWI groups for **Figure 7**. Pixels were assigned EBAL weights such that each SWI stratum had the same distribution of TPI, slope, ts-RBR and |

| | | | | |
|---|---|---|---|---|
| | | | | distance from nearest forest patch |
| 2.6 | Kaplan-Meier survival function | Stratified slope | Burnt grassland to young shrubland succession rate<br><br>Burnt shrubland to forest succession rate | Stratified into 20 slope groups for **Figure 7**. Pixels were assigned EBAL weights such that each slope stratum had the same distribution of SWI, ts-RBR and distance from nearest forest patch |
| 2.6 | Kaplan-Meier survival function | Stratified aspect | Burnt grassland to young shrubland succession rate<br><br>Burnt shrubland to forest succession rate | Stratified into 20 aspect groups for **Figure 7**. Pixels were assigned EBAL weights such that each aspect stratum had the same distribution of ts-RBR and distance from nearest forest patch |
| 2.6 | Kaplan-Meier survival function | Stratified ts-RBR | Burnt grassland to young shrubland succession rate<br><br>Burnt shrubland to forest succession rate | Stratified into five ts-RBR groups for **Figure S5c/S5d** and 20 ts-RBR groups for **Figure 7**. Pixels were assigned EBAL weights such that each ts-RBR stratum had the same distribution of TPI, SWI, slope, cos_aspect, and distance from nearest forest patch |
| 2.6 | Kaplan-Meier survival function | Stratified distance from nearest forest patch | Burnt grassland to young shrubland succession rate<br><br>Burnt shrubland to forest succession rate | Stratified into five distance groups for **Figure S5a/S5b** and 20 distance groups for **Figure 7**. Pixels were assigned EBAL weights such that each ts-distance stratum had the same distribution of TPI, SWI, slope, cos_aspect, and ts-RBR |
| 2.6 | Random survival forest | Prefire vegetation type, post-fire distance to nearest forest patch, ts-RBR, TPI, SWI, cos_aspect, slope | Expected time it takes (years) for burnt pixels to recover to forests | For assessing the importance of different variables in predicting post-fire recovery rate. Results were plotted in **Figure 6** |

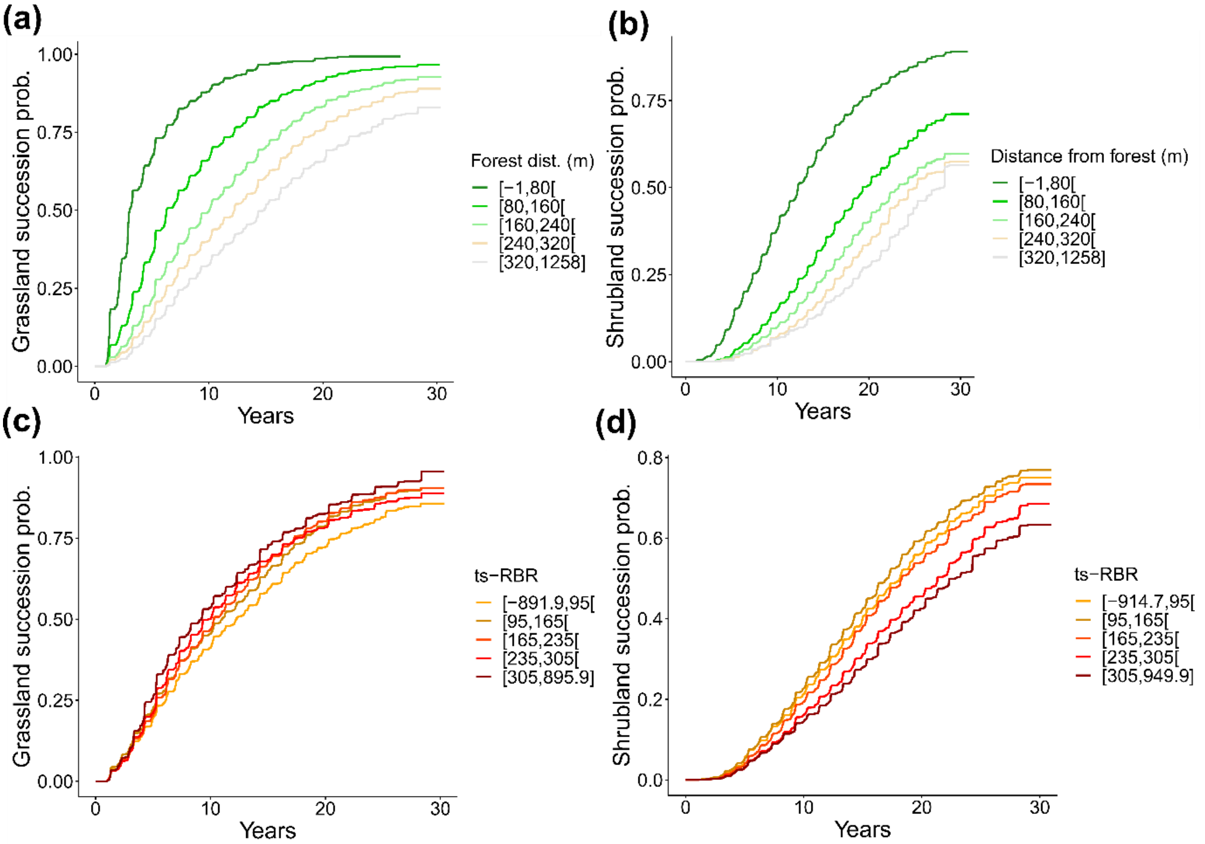


Figure S5: Kaplan-Meier survival curves built from a dataset stratified by distance to the nearest forest patch (a and b) or by burn severity (c and d). Panels (a) and (c) represent burnt grassland to shrubland succession probability over time, while panels (b) and (d) represent burnt shrubland to forest succession probability over time.
